# Assembly and lipid-gating of LRRC8A:D volume-regulated anion channels

**DOI:** 10.1101/2024.11.24.625074

**Authors:** Antony Lurie, David M. Kern, Katharine Henn, Stephen G. Brohawn

**Affiliations:** Department of Molecular & Cell Biology, University of California, Berkeley, CA, USA; Department of Neuroscience, University of California, Berkeley, CA, USA; California Institute for Quantitative Biology (QB3), University of California, Berkeley, CA, USA; Department of Chemistry, University of California, Berkeley, CA, USA

## Abstract

Volume-regulated anion channels (VRACs) are ubiquitously expressed vertebrate ion channels that open in response to hypotonic swelling. VRACs assemble as heteromers of LRRC8A and LRRC8B-E subunits, with different subunit combinations resulting in channels with different properties. Recent studies have described the structures of LRRC8A:C VRACs, but how other VRACs assemble, and which structural features are conserved or variant across channel assemblies remains unknown. Herein, we used cryo-EM to determine structures of a LRRC8A:D VRAC with a 4:2 subunit stoichiometry, which we captured in two conformations. The presence of LRRC8D subunits increases hydrophobicity and widens the selectivity filter, which may explain the unique substrate selectivity of LRRC8D-containing VRACs. The structures reveal lipids bound inside the channel pore, similar to those observed in LRRC8A:C VRACs. Using electrophysiological experiments, we confirmed that pore lipids block conduction in the closed state, demonstrating that lipid-gating is a general property of VRACs. Finally, we observe that LRRC8D subunit incorporation disrupts packing of the cytoplasmic LRR domains, increasing channel dynamics and opening lateral fenestrations, which we speculate are necessary for pore lipid evacuation and channel activation.

## Introduction

Volume-regulated anion channels (VRACs) are a family of ubiquitously expressed osmosensitive vertebrate ion channels that mediate cellular volume regulation and paracrine signaling^1,2^. VRACs are hexameric large pore channels which assemble as heteromers of LRRC8A and LRRC8B–E subunits^3–5^. With up to 1,934 unique channel assemblies possible, VRAC assembly diversity is thought to account for the observed variability in channel properties, including in single channel conductance, open probability, rectification, voltage-dependent inactivation, and substrate selectivity^2,3,5–7^. How differences in channel composition are structurally manifested to modulate channel properties remains largely unknown.

Recently, studies from our group^8^ and Rutz *et al.*^9^ described the first high-resolution structures of heteromeric LRRC8A:C VRACs. Both studies identified a broadly similar hexameric channel architecture, but with different LRRC8A:C stoichiometries of 5:1 or 4:2, respectively. We additionally observed lipids bound inside the pore which occlude the channel in the closed state. How other heteromeric LRRC8 channels assemble and whether they are similarly lipid-gated is unknown. Compared to other characterized VRAC assemblies (i.e. LRRC8A:C/E), LRRC8A:D VRACs show smaller Cl^−^ conductances, greater open probabilities, and increased permeation of large and non-anionic substrates including taurine, γ-aminobutyric acid (GABA), *myo*-inositol, cisplatin, and blasticidin^3,5–7,10^. By determining cryo-EM structures of a LRRC8A:D VRAC, we aimed to identify which features are shared with LRRC8A:C VRACs, while also establishing a structural basis for their unique channel properties.

## Results and Discussion

To study LRRC8A:D VRACs by cryo-EM, we leveraged our previously described approach for characterizing fiducially-tagged LRRC8A:C VRACs (Extended Data Figure 1)^8^. Mouse LRRC8A and LRRC8D subunits were over-expressed in insect cells (which lack endogenous LRRC8s) using a single engineered baculovirus. Each subunit was C-terminally tagged with a fluorescent protein fused through a protease-cleavable linker, with LRRC8A and LRRC8D subunits bearing different fluorescent proteins and protease cleavage tags to enable subunit-specific pulldown. To eliminate channel pseudosymmetry for cryo-EM studies, a BRIL domain^11,12^ was inserted into the extracellular loop of LRRC8A. Fiducial mass was further increased by adding an α-BRIL antibody fragment (Fab)^13^ and an α-Fab nanobody (Nb)^14^ to the purified sample. Detergent solubilization, followed by two sequential rounds of affinity chromatography, allowed us to specifically isolate LRRC8A:D VRACs for cryo-EM single particle analysis.

We initially resolved a consensus map at an overall resolution of 3.1 Å (Extended Data Figure 2), revealing a hexameric LRRC8 channel. However, poor map quality in three of the six subunits precluded confident assignment of each subunit’s identity. Using 3D variability analysis in cryoSPARC^15^, we resolved two conformations with improved local map qualities to overall resolutions of 3.3 – 3.4 Å. These reconstructions allowed us to assign identities to all subunits and to construct atomic models incorporating the transmembrane domains, extracellular regions, and linker regions of all subunits (Figure 1A – B, Extended Data Figures 2 – 3). On the basis of fiducial and subunit-specific side chain densities (Extended Data Figure 4), we inferred that both conformations correspond to the same channel assembly with a 4:2 LRRC8A:D stoichiometry and adjacent LRRC8D subunits. Our interpretation of these data is that the 4:2 LRRC8A:D assembly is the most prevalent in our purified sample. It is possible that additional assemblies are present, but are too structurally heterogenous or rare to be resolved. Unless otherwise noted, we focus our analyses on the better resolved conformation 1.

**Figure 1.**
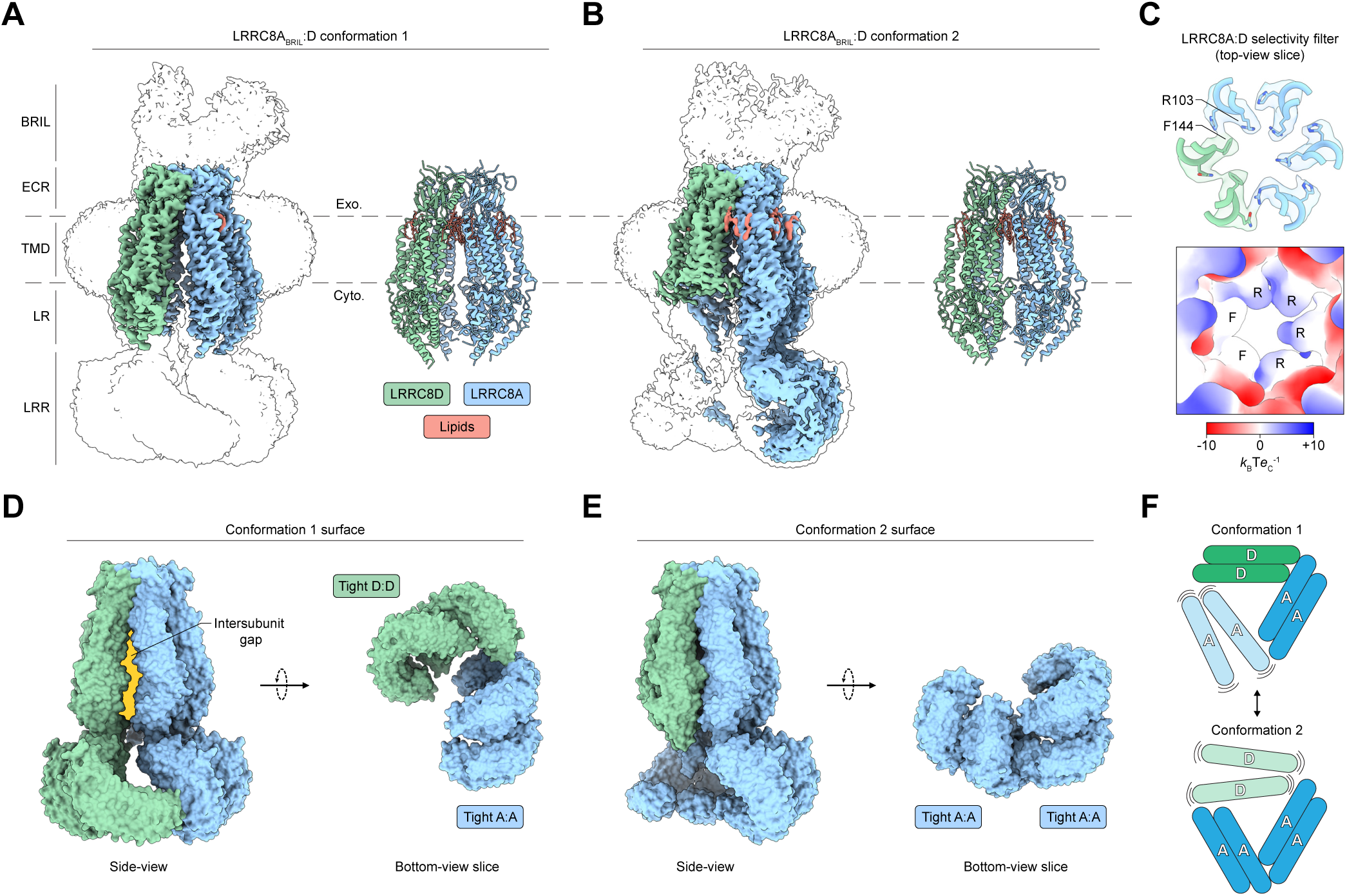
Architecture of a LRRC8A_BRIL_:D VRAC. **A – B)** Side-views of cryo-EM density maps (*left*) and their associated models (*right*) for LRRC8A:D conformation 1 (**A**) and 2 (**B**). **C)** Focused top-view of the LRRC8A:D selectivity filter model overlaid with its carved density (*top*) and the model surface colored by electrostatic potential (*bottom*). **D – E)** Side-view (*left*) and bottom-view slice (*right*) of the LRRC8A:D surface with docked LRRs for LRRC8A:D conformation 1 (**D**) and 2 (**E**). The intersubunit gap induced by ordering of the LRRC8D LRR pair in conformation 1 (**D**, *left*) is highlighted in yellow. **F)** Cartoon schematic of the LRRC8A:D LRR conformations. LRRC8A subunits, blue; LRRC8D subunits, green; lipids, salmon. Cyto, cytoplasmic; ECR, extracellular region; Exo, exoplasmic; LR, linker region; TMD, transmembrane domain.

Comparing LRRC8A:D to previously described structures of LRRC8A:C^8^, LRRC8A homomers^16^, and LRRC8D homomers^17^ shows that LRRC8A:D follows a similar overall architecture with a nearly invariant extracellular region (pairwise backbone r.m.s.d. < 1 Å). Within this rigid extracellular scaffold, differences in selectivity filter residues may explain differences in substrate selectivity between channel assemblies. The selectivity filter is formed by a single residue from each subunit pointing towards the conduction axis: R103 for LRRC8A, L105 for LRRC8C, and F144 for LRRC8D. LRRC8D subunit incorporation alters both the size and chemistry of the filter. Introduction of F144 residues from LRRC8D subunits increases hydrophobicity along one face of the filter compared to LRRC8A homomeric channels (Figure 1C). The F144 side chain also permits more dilated filters since it lays flatter against the outer vestibule wall compared to L105 in LRRC8C or R103 in LRRC8A (Figure 2). While the structures of LRRC8A:D presented here are similarly constricted at the selectivity filter as LRRC8A:C (to ∼5.4 Å in diameter for LRRC8A:D and ∼5.5 Å for LRRC8A:C), this is due to an extended rotamer adopted by one LRRC8A R103 residue. We predicted that assemblies with higher LRRC8D to LRRC8A subunit ratios would house expanded selectivity filters. Indeed, a model of a 1:5 LRRC8A:D selectivity filter shows a predicted minimum pore diameter of ∼7.6 Å, only slightly smaller than that observed in a structure of a non-physiological LRRC8D homomer (∼8.2 Å). The expanded size and increased hydrophobicity of LRRC8D-containing channel filters thus likely underlies their greater permeability to non-anionic or larger substrates.

**Figure 2.**
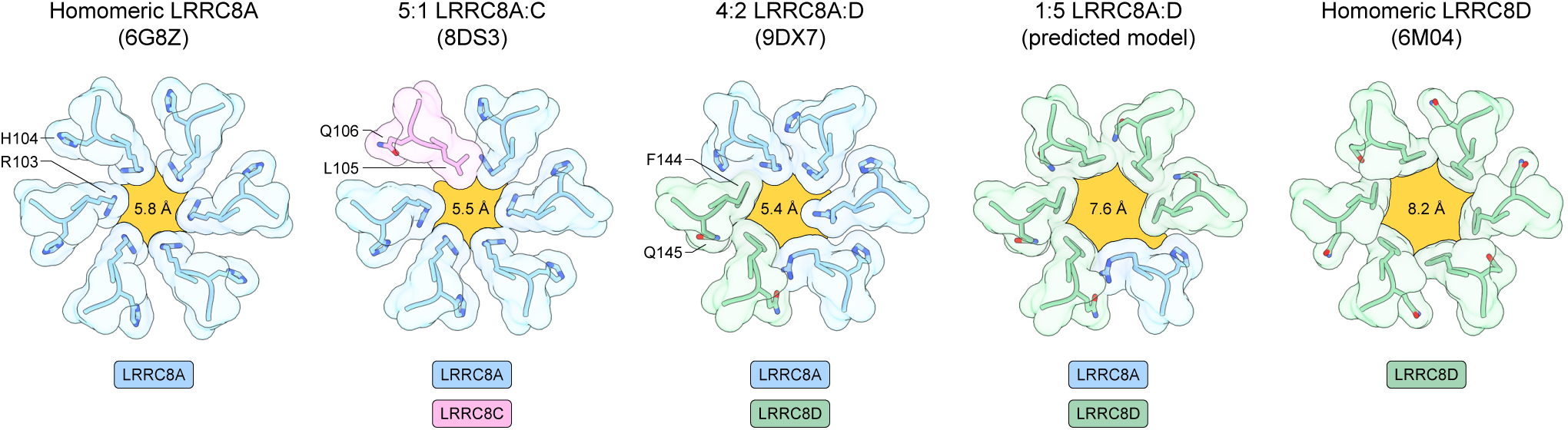
Comparison of VRAC selectivity filters. Focused top-views of selected selectivity filter models. The pore is outlined and highlighted in yellow and overlaid with the calculated minimum pore diameter. LRRC8A subunits, blue; LRRC8C subunits, pink; LRRC8D subunits, green.

The major structural differences between LRRC8A:D conformations and other VRAC structures arise from the cytoplasmic leucine-rich repeat domains (LRRs). Although poor local resolutions precluded their atomic modeling, the positions of four LRRs were sufficiently resolved to permit docking of LRR models for visualization purposes (Figure 1D – E). Shared across both conformations, one pair of LRRC8A LRRs adopts an ordered “tight” interaction identical to those observed in structures of LRRC8A homomers^16,18,19^ and LRRC8A:C heteromers^8,9^. In conformation 1, two LRRC8D subunits form a second tight LRR pair, similar to those observed in a homomeric LRRC8D structure^17^, while the two remaining LRRC8A LRRs are disordered (Figure 1D). The opposite is true in conformation 2, where the LRRC8D LRRs are disordered, but the two remaining LRRC8A subunits order to form a second tight LRR pair (Figure 1E). These LRR dynamics can be conceptualized as a bistable equilibrium, with stabilization of a LRRC8A LRR pair in competition with, and alternating, with stabilization of the LRRC8D LRR pair (Figure 1F). As in LRRC8A:C^8,9^, incorporation of non-LRRC8A (i.e. LRRC8D) subunits prevents the LRRs from adopting the stable pseudo-symmetric trimer of dimers arrangement seen in structures of LRRC8A homomers^16,18,19^. Here, only four LRRs can adopt a stable position at a time.

Destabilization of the LRR arrangement by LRRC8D subunits propagates up the channel to alter the positions of linker regions and transmembrane domains. These changes are stark in conformation 1, where ordering of the LRRC8D LRR pair causes one LRRC8D subunit to peel away from the conduction axis (Figure 1D). This creates a large lateral intersubunit fenestration open to ∼6 Å or wider at the inner leaflet of the membrane; wide enough to permit passage of lipids into and out of the channel pore. Notably, our recent structures of a LRRC8A:C VRAC similarly found that incorporation of a LRRC8C subunit results in increased LRR dynamics and opening of a lateral fenestration capable of passing lipids, suggesting that these properties may be necessary for channel function^8^. One possibility is that lateral fenestrations stabilized by heterotypic LRR interactions may allow for the removal of pore-occluding lipids (discussed below), which would be impeded in a rigid channel closed off from the membrane. This may help explain why incorporation of non-LRRC8A subunits is necessary for full channel activation^3,5^ as LRRC8A homomeric channels show reduced LRR dynamics, maintain small intersubunit gaps, and generate only small currents in cells.

We observe density inside the LRRC8A:D channel pore consistent with lipids that occlude the conduction axis, reminiscent of those found in LRRC8A:C and chimeric LRRC8A-C structures (Figure 3)^8,20^. The presence of pore lipids constricts the pore to a radius of 1.6 – 1.9 Å and is expected to result in channel closure (Figure 3C). To test whether the observed pore lipids contribute to channel gating in LRRC8A:D VRACs, we used whole-cell patch clamp electrophysiology to characterize hypotonicity-induced currents from wild-type and mutant LRRC8A:D channels expressed in *LRRC8A-E^−/–^* HeLa cells (Figure 4, Extended Data Figure 5). In cells transfected with wild-type LRRC8A and LRRC8D subunits, we observed robust channel activation (mean fold activation ± s.e.m. = 20.9 ± 3.9) upon hypotonicity-induced cell swelling, consistent with wild-type LRRC8A:D currents. To examine the role of pore lipids in channel gating, we introduced the T48D mutation into LRRC8A and/or LRRC8D subunits, which is predicted to electrostatically impede lipid occupancy in the channel pore and was found to result in constitutive channel activity in LRRC8A:C VRACs (Figure 4A)^8^. Introduction of T48D into LRRC8D alone was not sufficient to induce significant constitutive channel activity (28.8 ± 4.6, n.s.). However, introduction of T48D into LRRC8A resulted in constitutively active channels with reduced fold activation upon swelling (2.8 ± 0.4, *P* = 0.0050 compared to WT). Incorporation of the T48D mutation in both LRRC8A and LRRC8D subunits further reduced fold activation (1.6 ± 0.2, *P* = 0.0032 compared to WT, *P* = 0.0326 compared to LRRC8A(T48D):D). The increased penetrance of the T48D mutation in LRRC8A compared to LRRC8D is consistent with LRRC8A:D VRACs containing more LRRC8A than LRRC8D subunits under these expression conditions. We conclude that LRRC8A:D VRACs are constricted in the closed state by pore lipids.

**Figure 3.**
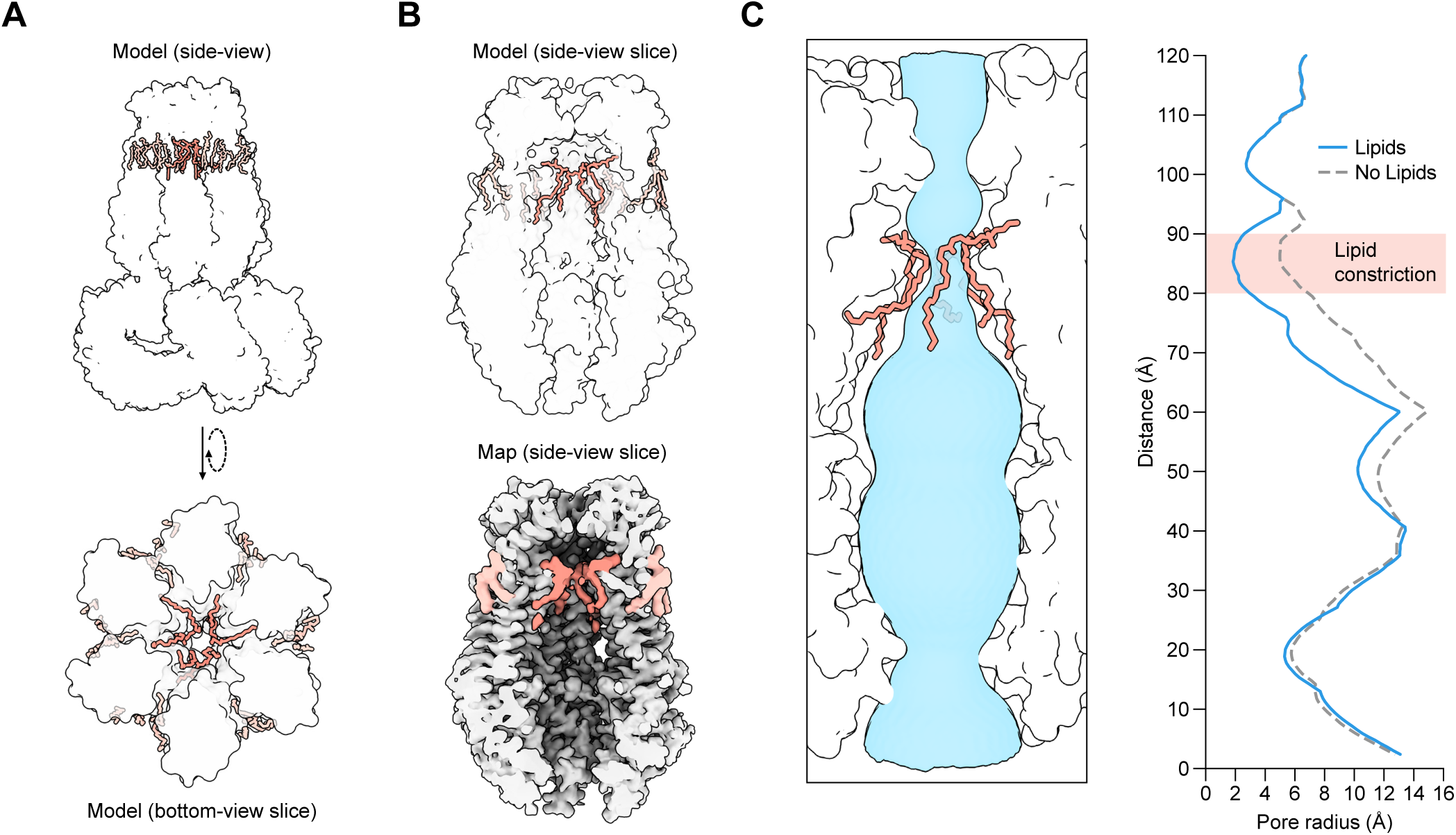
LRRC8A:D VRACs are closed by pore lipids. **A)** Side-view (*top*) and bottom-view slice (*bottom*) of the LRRC8A:D model with pore (salmon, darker) and annular (salmon, lighter) lipids highlighted. **B)** Side-view slices of the LRRC8A:D model (*top*) and carved map density (*bottom*) with pore and annular lipids highlighted. **C)** *Left*: side-view slice of the LRRC8A:D model pore with lipids and the pore surface (blue) highlighted. *Right:* Calculated pore radii in the presence (blue, solid line) and absence (gray, dashed line) of pore lipids.

**Figure 4.**
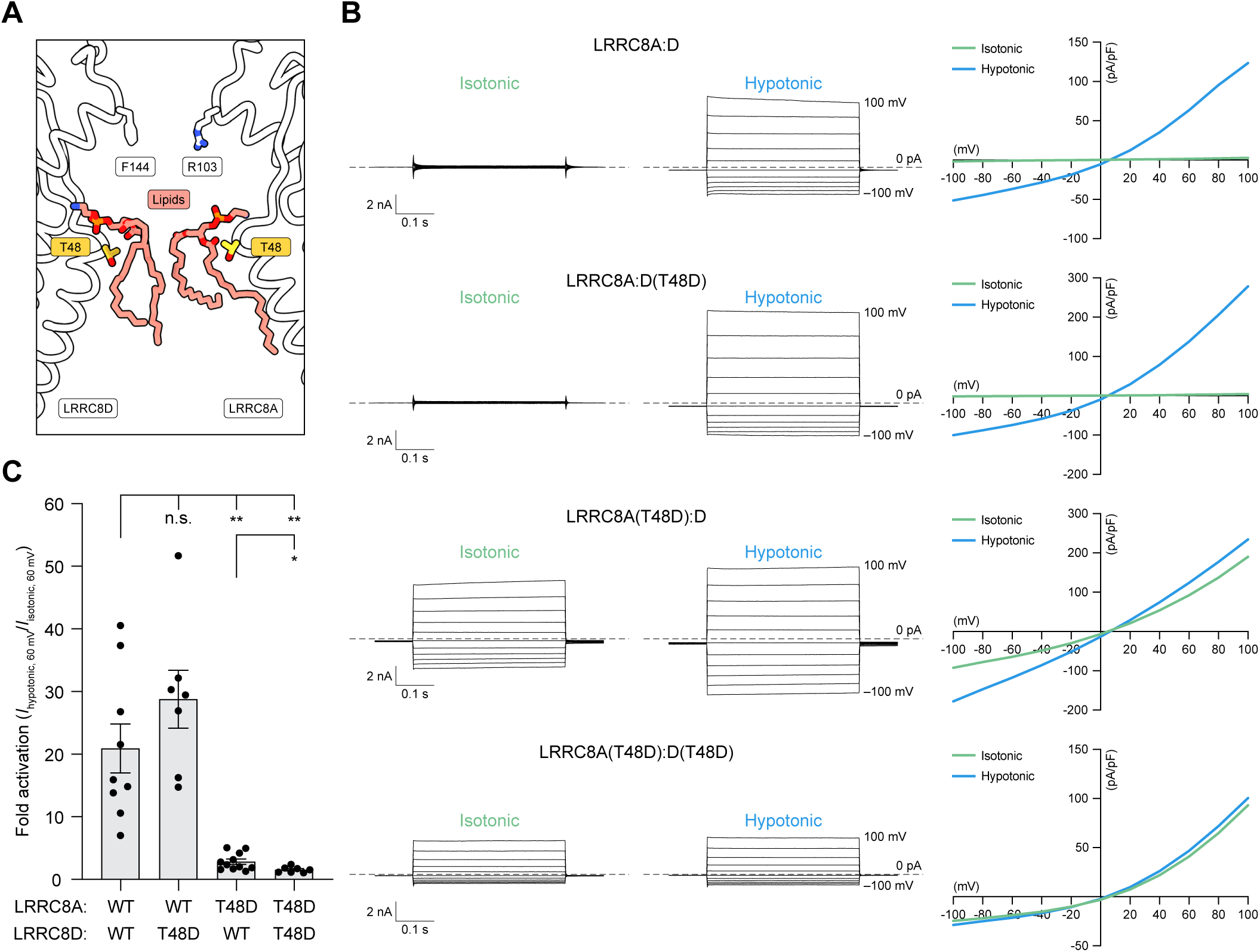
LRRC8A:D VRACs are gated open upon removal of pore lipids. **A)** Focused side-view slice of the LRRC8A:D model pore with selectivity filter residues (white), pore lipids (salmon), and residue T48 (yellow) highlighted. **B)** Whole-cell voltage-clamp recordings from *LRRC8A-E^−/–^* HeLa cells expressing wild-type (WT) LRRC8A:D (*top*), LRRC8A:D(T48D) (*center-top*), LRRCA(T48D):D (*center-bottom*), or LRRC8A(T48D):D(T48D) (*bottom*) mutant channels. For each construct, representative current traces from isotonic (*left*) and hypotonic (*center*) solutions are displayed alongside corresponding plots of the current-voltage relationships (*right*; isotonic, green; hypotonic, blue). For current traces, 0 pA/pF is marked with a dotted line. **C)** Fold-activation of WT and mutant LRRC8A:D channels following hypotonic swelling (*I*_hypotonic, 60 mV_ / *I*_isotonic, 60 mV_). Data are displayed as the mean ± s.e.m. plotted alongside individual data points for WT LRRC8A:D (*n* = 9), LRRC8A:D(T48D) (*n* = 7), LRRC8A(T48D):D (*n* = 11), and LRRC8A(T48D):D(T48D) (*n* = 7). Differences were assessed using a Brown-Forsythe and Welch one-way ANOVA test followed by a Dunnett’s T3 multiple comparisons test to WT LRRC8A:D (n.s., not significant; ** *P* = 0.0050 and *P* = 0.0032 for LRRC8A(T48D):D and LRRC8A(T48D):D(T48D), respectively) and an unpaired *t* test comparing LRRC8A(T48D):D to LRRC8A(T48D):D(T48D) (* *P* = 0.0326).

In conclusion, we resolved two conformations of a LRRC8A:D VRAC, revealing a hexameric assembly with high structural variability in the cytoplasmic LRR domains and linker regions. These structures show how incorporation of LRRC8D subunits increases hydrophobicity and expands the selectivity filter, which may underly the unique selectivity of LRRC8D-containing VRACs for large and non-anionic substrates. On the other hand, the structural determinants of other channel properties, including single channel conductance and open probability, remain unknown. We also observe lipids bound within the channel pore, similar to those observed in LRRC8A:C VRACs, and validated their role in channel gating using electrophysiology. We thus expect the pore lipid gate to be present in all VRAC assemblies and that lipid-gating is a general property of these channels. Finally, we found that heterotypic LRR interactions destabilize the channel’s structure to pry open lateral fenestrations capable of lipid permeation. We speculate that these fenestrations form paths for evacuating pore lipids and may thus be necessary for channel opening.

---

**Extended Data Figure 1.**
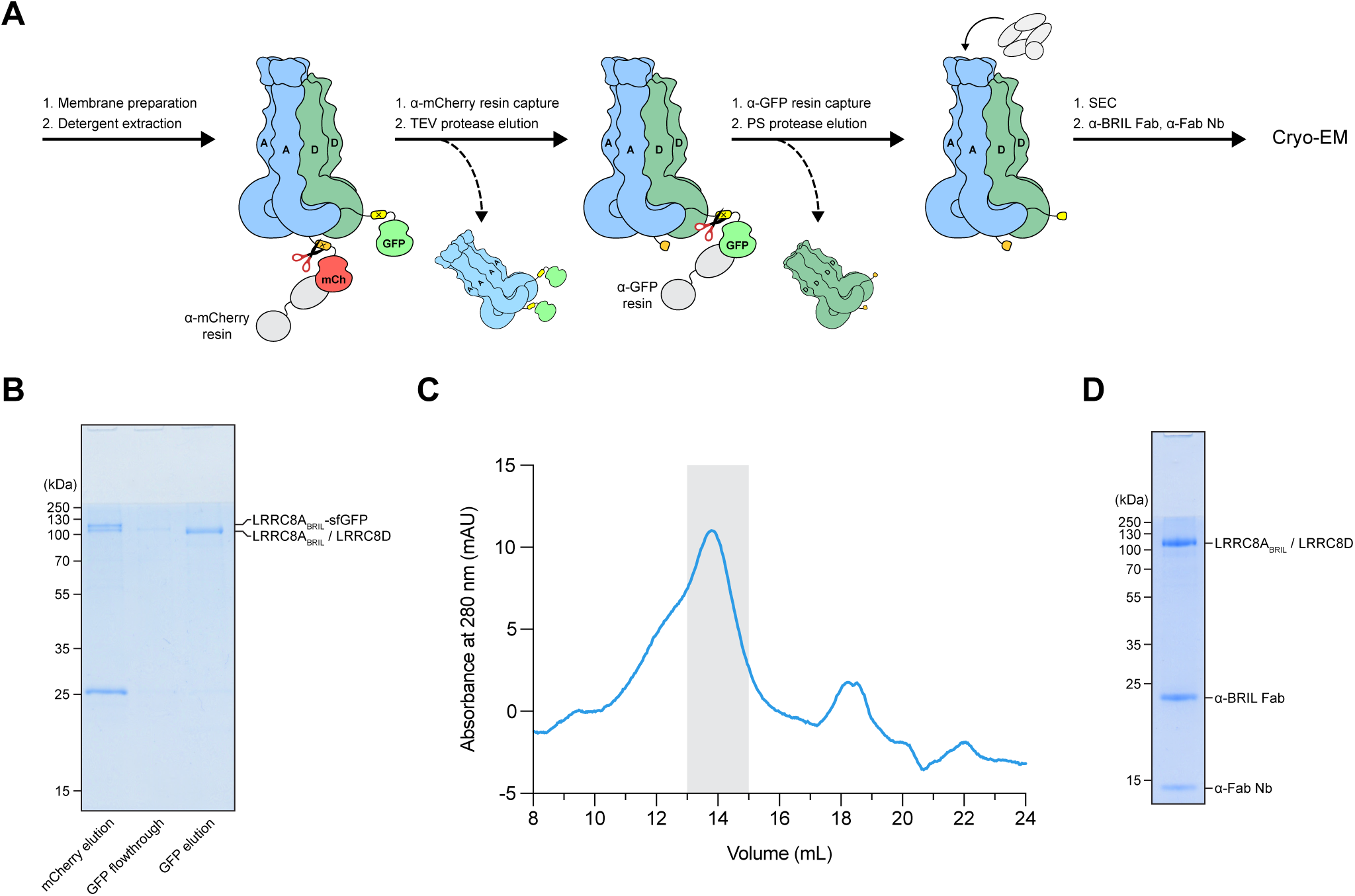
Purification of LRRC8A_BRIL_:D VRACs. **A)** Schematic of the LRRC8A_BRIL_:D purification approach. **B)** Coomassie-stained SDS-PAGE of LRRC8A_BRIL_:D purification samples following elution from the mCherry Nb resin (*left*), flow-through after GFP Nb resin binding (*center*), and elution from the GFP Nb resin (*right*). **C)** Size-exclusion chromatogram of the crude LRRC8A_BRIL_:D sample (blue line, monitoring by absorbance at 280 nm) with pooled sample fractions highlighted in gray. **D)** Coomassie-stained SDS-PAGE of the final LRRC8A_BRIL_:D sample with added α-BRIL Fab (BAG2) and α-Fab Nb. mCh, mCherry; PS, PreScission; SEC, size-exclusion chromatography.

**Extended Data Figure 2.**
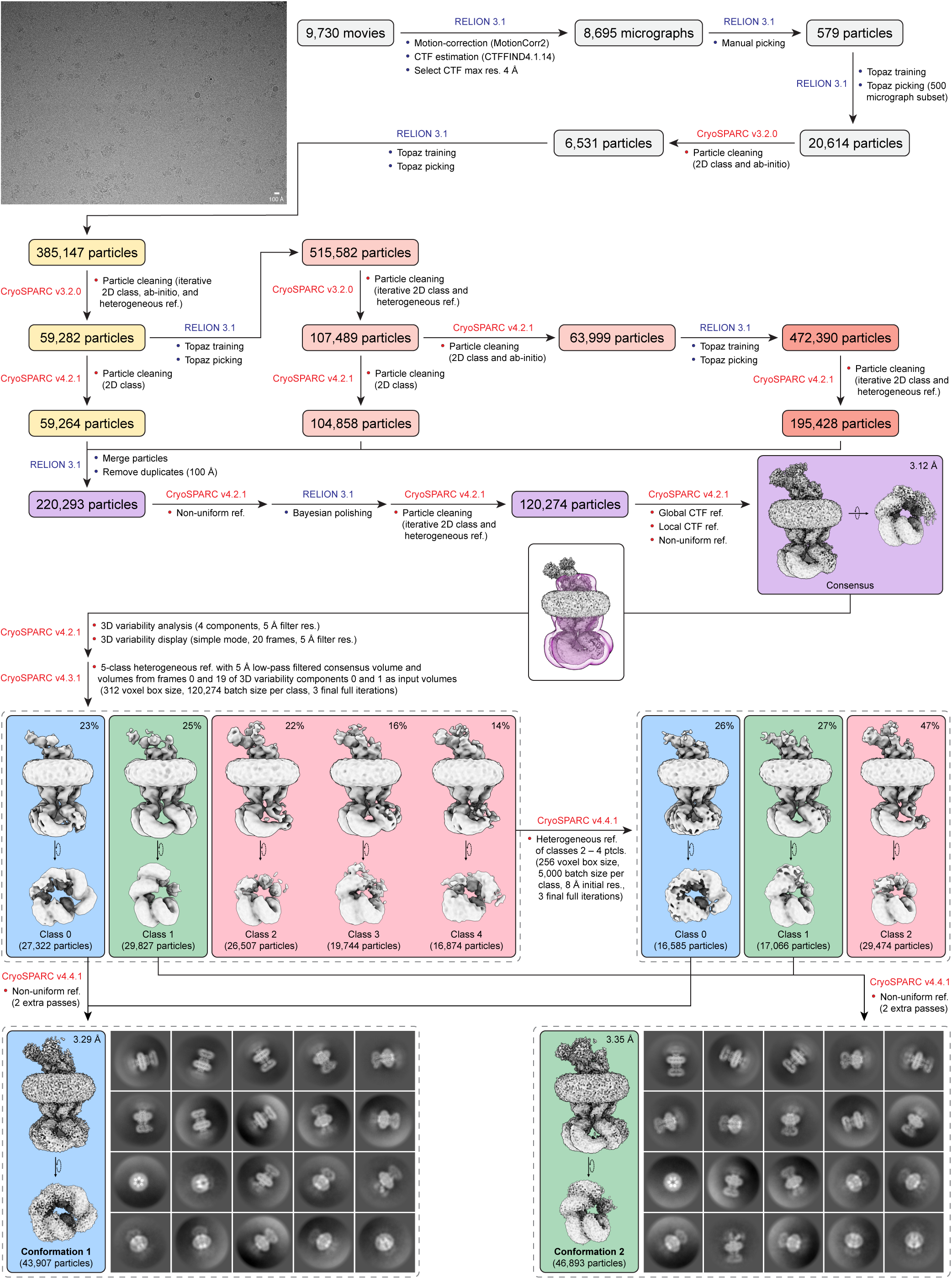
Summarized workflow for cryo-EM data processing. A representative micrograph is displayed in the top left corner. Representative 2D classes from the final particle stacks are displayed on the bottom.

**Extended Data Figure 3.**
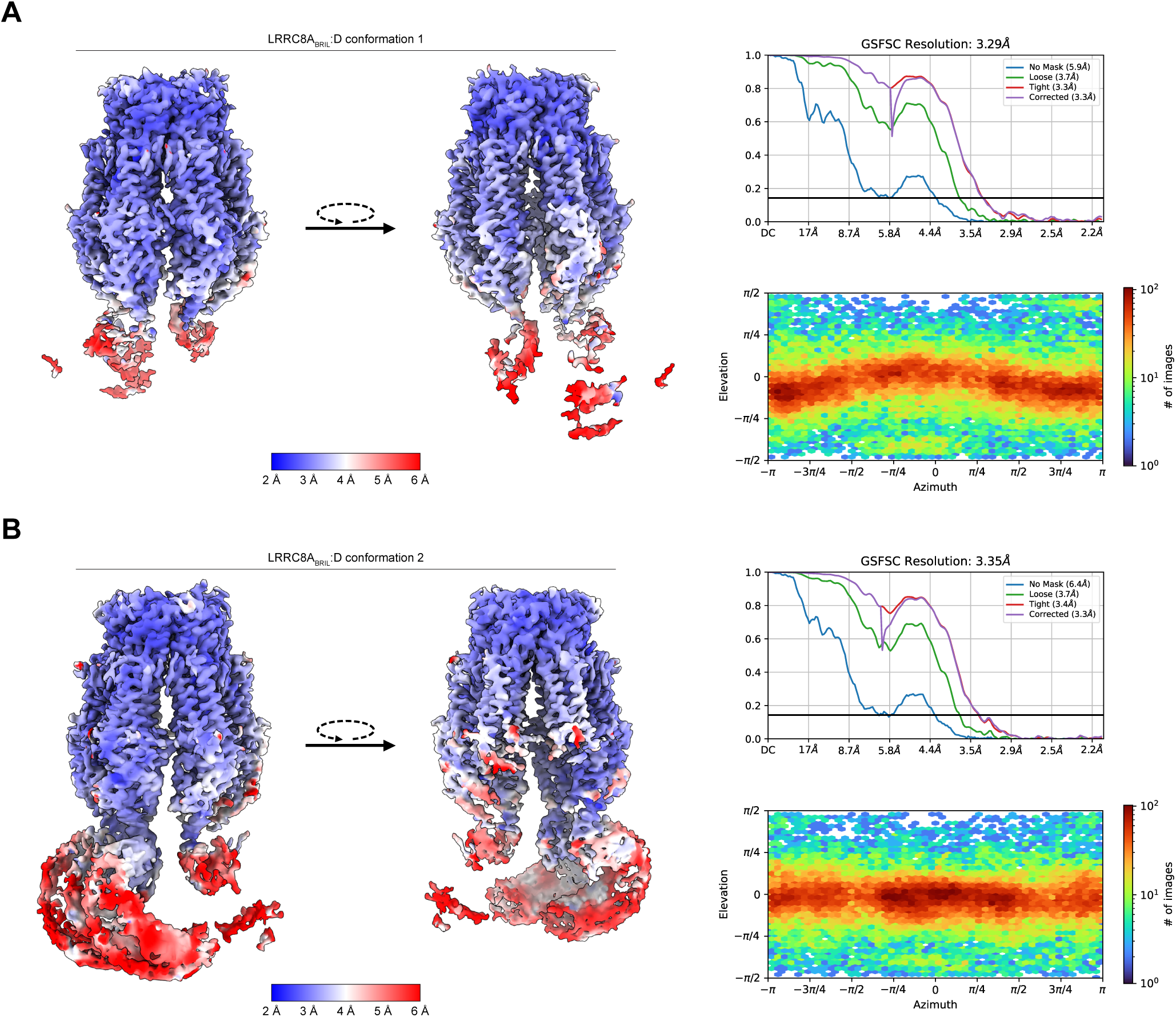
Validation of cryo-EM data. Two side views of map density colored by local resolution (*left*), FSC plots (*right, top*), and viewing direction distribution plots (*right, bottom*) for LRRC8A:D conformation 1 (**A**) and 2 (**B**).

**Extended Data Figure 4.**
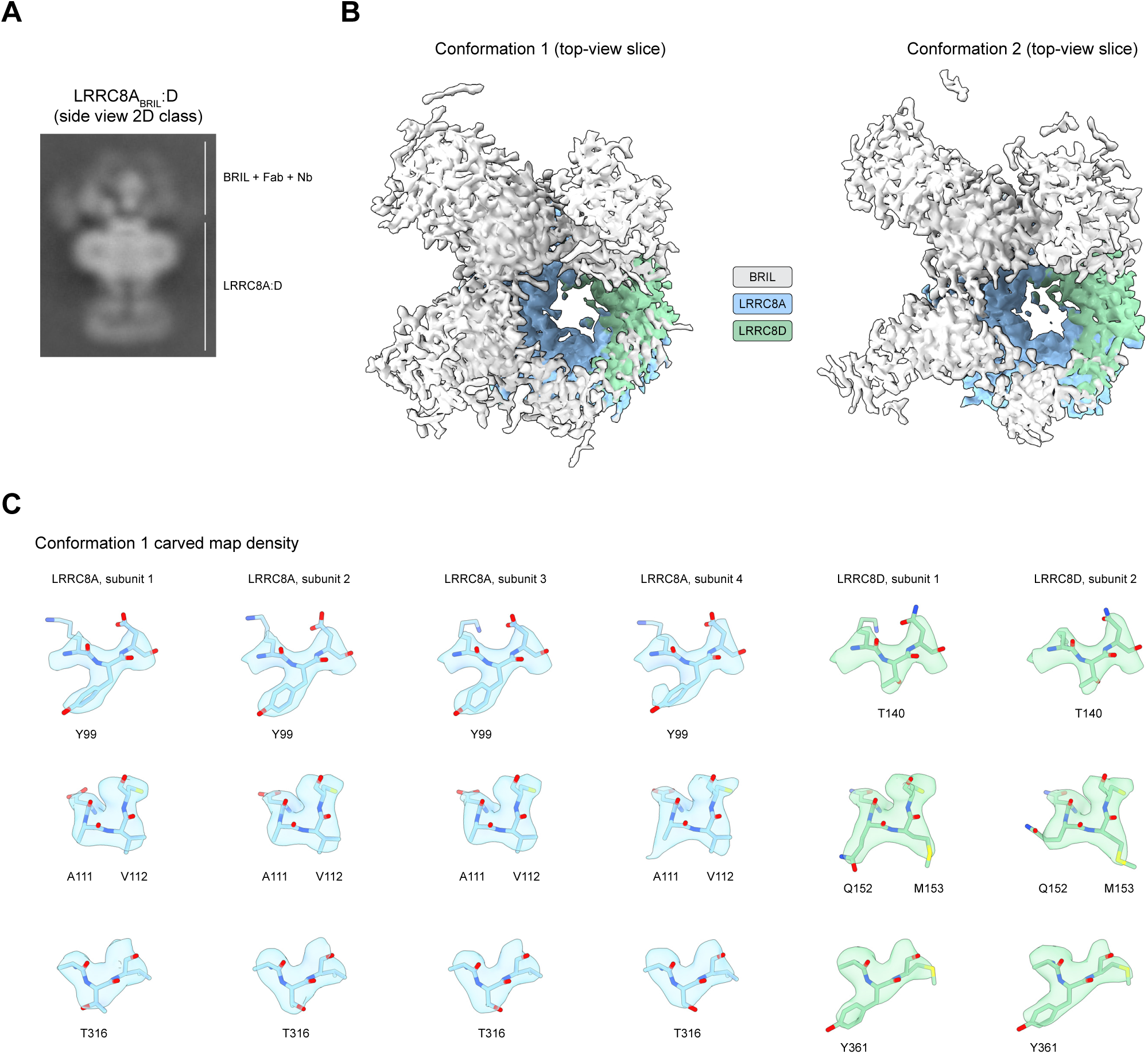
Validation of subunit assignment. **A)** 2D class of LRRC8A_BRIL_:D exhibiting density for BRIL / α-BRIL Fab / α-Fab Nb complexes. **B)** Top-view slices of the LRRC8A:D conformation 1 (*left*) and 2 (*right*) maps focusing on the extracellular region (LRRC8A_BRIL_, blue; LRRC8D, green) and globular BRIL densities (gray), highlighting density for a BRIL domain positioned above each of the four LRRC8A_BRIL_ subunits. **C)** Selected residues with sequence differences between LRRC8A and LRRC8D overlaid with their carved cryo-EM map densities. The residue comparisons displayed are: Y99 (LRRC8A) to T140 (LRRC8D), A111–V112 (LRRC8A) to Q152–M153 (LRRC8D), and T316 (LRRC8A) to Y361 (LRRC8D).

**Extended Data Figure 5.**
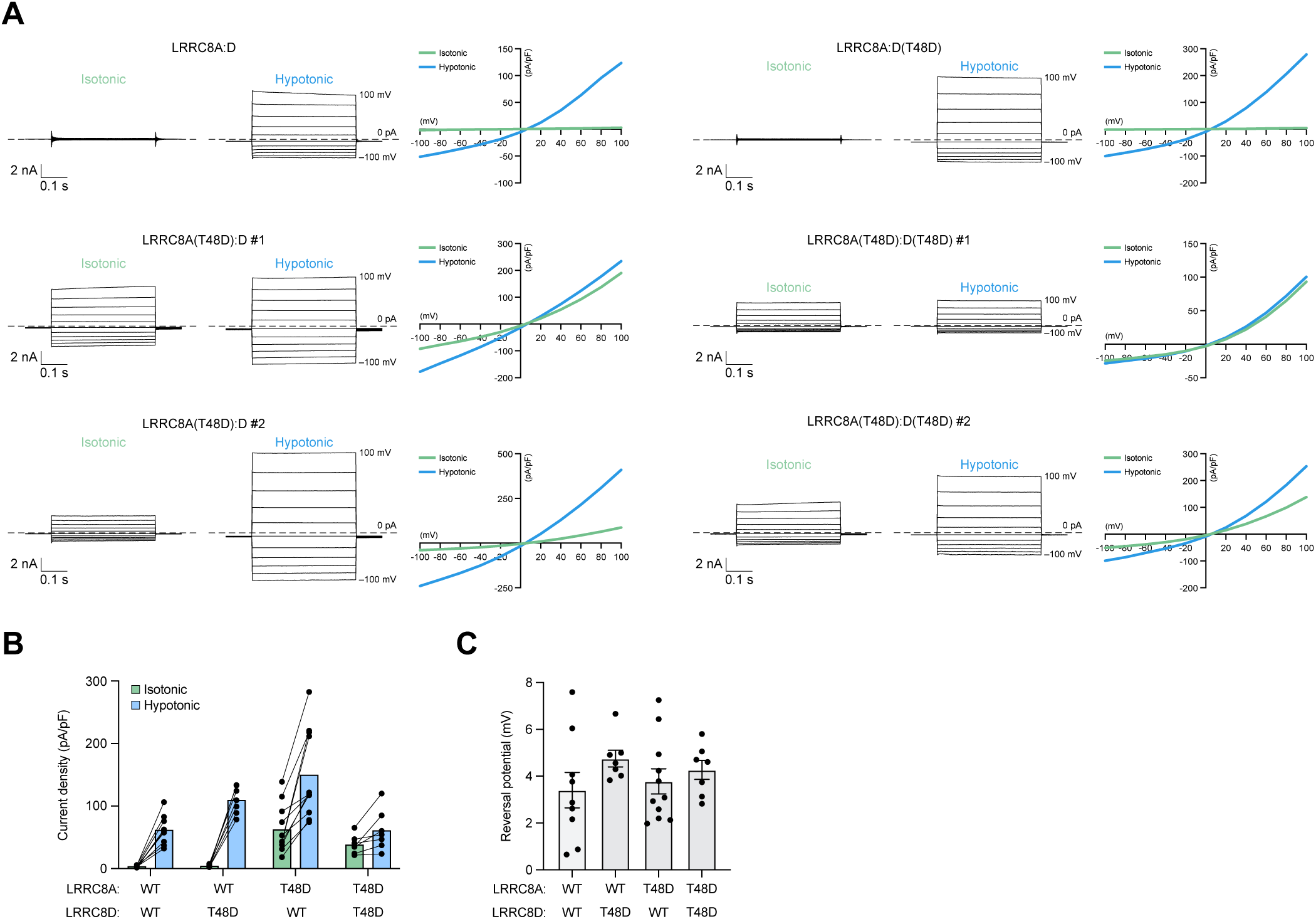
Electrophysiology of LRRC8A:D lipid gate mutants. **A)** Whole-cell voltage-clamp recordings from *LRRC8A-E^−/–^* HeLa cells expressing wild-type (WT) LRRC8A:D, LRRC8A:D(T48D), LRRCA(T48D):D, or LRRC8A(T48D):D(T48D) mutant channels. For each, representative current traces from isotonic (*left*) and hypotonic (*center*) solutions are displayed alongside corresponding plots of the current-voltage relationships (*right*; isotonic, green; hypotonic, blue). Due to variability in the observed fold activation, two representative examples are shown for LRRCA(T48D):D and LRRC8A(T48D):D(T48D). For current traces, 0 pA/pF is marked with a dotted line. **B)** Current densities for WT LRRC8A:D (*n* = 9), LRRC8A:D(T48D) (*n* = 7), LRRC8A(T48D):D (*n* = 11), and LRRC8A(T48D):D(T48D) (*n* = 7) channels in isotonic (green) and hypotonic (blue) solutions. Data are displayed as the mean plotted alongside paired individual data points for each cell. **C)** Reversal potentials for WT LRRC8A:D (*n* = 9), LRRC8A:D(T48D) (*n* = 7), LRRC8A(T48D):D (*n* = 11), and LRRC8A(T48D):D(T48D) (*n* = 7) channels following hypotonic swelling.

## Methods

### Construct design and protein expression

The protein expression construct for LRRC8A_BRIL_ was described previously^8^. Briefly, the *Mus musculus* LRRC8A sequence (UniProt: Q80WG5) was codon optimized for *Spodoptera frugiperda* and cloned into a custom vector based on the pACEBAC1 backbone (MultiBac; Geneva Biotech, Geneva, Switzerland) with an added C-terminal PreScission protease cleavage site, linker sequence, superfolder GFP (sfGFP) and 7xHis tag. The sequence for apocytochrome b562RIL (BRIL)^11,21^ was then codon optimized for *Spodoptera frugiperda* and inserted into the LRRC8A extracellular loop, between residues 76 and 91, generating a construct for expression of mmLRRC8A(76-BRIL-91)-SNS-LEVLFQGP-SRGGSGAAAGSGSGS-sfGFP-GSS-7xHis (LRRC8A_BRIL_). The coding sequence for *Mus musculus* LRRC8D (UniProt: Q8BGR2) was codon optimized for *Spodoptera frugiperda*, synthesized (Gen9, Cambridge, MA), and cloned into a custom vector based on the pACEBAC1 backbone (MultiBac) with an added C-terminal TEV protease cleavage site, linker sequence, and mCherry tag, generating a construct for expression of mmLRRC8D-SNS-ENLYFQG-SRGSGSGS-mCherry. The LRRC8A and LRRC8D cassettes, each with a polyhedrin promoter and SV40 terminator, were iteratively cloned using the I-CeuI and BstXI sites in the pACEBAC1 backbones to generate the dual LRRC8A_BRIL_:D expression plasmid. MultiBac cells were then used to generate a LRRC8A_BRIL_:D bacmid according to the manufacturer’s instructions.

For electrophysiology, untagged versions of *Mus musculus* LRRC8A and LRRC8D were each cloned into mammalian expression vectors containing a cytomegalovirus (CMV) promoter and IRES eGFP or mCherry expression reporters, respectively. The LRRC8A(T48D) mutant construct was described previously^8^. The T48D mutation was introduced into LRRC8D by PCR using the following primers: GATGCAACTGGATAAGGACCAGG (forward) and GTACCAGCGAATATAGC (reverse).

### Channel expression and purification

For protein production, Sf9 cells (Expression Systems, Davis, CA) were cultured in ESF 921 medium (Expression Systems) supplemented with 100 units/mL penicillin, 100 μg/mL streptomycin, and 0.25 μg/mL Amphotericin B (Gibco) and grown at 27 °C, shaking at 130 rpm. Baculovirus generation and amplification were conducted as described previously^8,22^. Briefly, P1 baculovirus was generated from cells transfected with LRRC8A_BRIL_:D bacmid using Escort IV reagent (MilliporeSigma, Burlington, MA) according to the manufacturer’s instructions. P2 virus was generated by infecting cells at 2 million cells/mL with P1 virus at a multiplicity of infection (MOI) of ∼0.1. Infection was monitored by fluorescence and cells were harvested at 72 hours. P3 virus was generated in a similar manner to expand the viral stock. P3 virus was used to infect Sf9 cells at 4 million cells/mL at a MOI of ∼2 – 5. At 72 hours, infected cells expressing LRRC8A_BRIL_:D were harvested by centrifugation at 2,500 x g for 10 minutes and frozen at −80°C. Cell pellets for purification came from two batches of 1 L cultures (total: 2 L culture, ∼28 mL cell pellet).

For protein purification (summarized in Extended Data Figure 1), cell pellets were thawed and resuspended in 100 mL of lysis buffer (50 mM HEPES, 150 mM KCl, 1 mM EDTA, pH 7.4). Protease inhibitors (final concentrations: E64 (1 µM), pepstatin A (1 µg/mL), soy trypsin inhibitor (10 µg/mL), benzamidine (1 mM), aprotinin (1 µg/mL), leupeptin (1 µg/mL), AEBSF (1mM), and PMSF (1 mM)) were added to the lysis buffer immediately before use. Benzonase (4 µl) was added after cell thaw. Cells were lysed by sonication and centrifuged at 150,000 x g for 1 hour at 4 °C. The supernatant was discarded and residual nucleic acid was removed from the top of the membrane pellet using D-PBS. Membrane pellets were scooped into a Dounce homogenizer and homogenized in 150 mL of extraction buffer (50 mM HEPES, 150 mM KCl, 1 mM EDTA, 1% (m/v) glyco-diosgenin (GDN: Anatrace, Maumee, OH), all protease inhibitors present in lysis buffer except PMSF, pH 7.4). The resulting membrane homogenate was gently stirred for 2 hours at 4 °C to extract membrane proteins, followed by centrifugation at 30,000 x g for 45 min at 4 °C. The supernatant, containing solubilized membrane proteins, was bound to 5 mL of pre-washed sepharose resin coupled to α-mCherry nanobody with gentle stirring for 1 hour at 4 °C. The resin was then collected in a column and washed with 10 mL of buffer 1 (20 mM HEPES, 150 mM KCl, 1 mM EDTA, 0.02% GDN, pH 7.4), 40 mL of buffer 2 (20 mM HEPES, 500 mM KCl, 1 mM EDTA, 0.02% GDN, pH 7.4), and 10 mL of buffer 1. The resin was then resuspended with ∼6 mL of buffer 1 containing 1.5 mM dithiothreitol (DTT) and 1 mg of TEV protease and nutated gently in the capped column overnight at 4°C. Cleaved LRRC8 complexes were eluted with an additional ∼7 mL of buffer 1. Eluate was then applied to a column containing 5 mL of sepharose resin coupled to α-GFP nanobody for a total of five passes through the resin. The resin was then washed with 20 mL of buffer 1, 10 mL of buffer 2, and another 10 mL of buffer 1. The resin was then resuspended with 6 mL of Buffer 1 containing 0.5 mg of PreScission protease and nutated gently in the capped column for 2 hours at 4 °C. Cleaved LRRC8A_BRIL_:D heteromeric complexes were then eluted with an additional ∼7 mL of buffer 1, spin concentrated to ∼500 µl with a 100 kDa molecular weight cutoff Amicon Ultra spin concentrator (Millipore), and then loaded onto a Superose 6 Increase column (GE Healthcare, Chicago, IL) equilibrated in buffer 1 on an NGC system (Bio-Rad, Hercules, CA) running Chromlab 6.0. Peak fractions containing LRRC8A_BRIL_:D complexes were pooled and spin concentrated to < 500 μL. Purified α-BRIL Fab BAG2 and α-Fab Nb^8,12,13^ were thawed, diluted with buffer 1, and added in molar excess to LRRC8A_BRIL_:D complexes at a ratio of ∼1:1.5:3 (LRRC8 : BAG2 : Nb) and gently nutated at 4 °C for 30 min. The mixture was spin concentrated, centrifuged at 21,000 x g for 5 minutes at 4 °C, and the supernatant (∼8 µL at 1.9 mg/mL) used immediately for cryo-EM sample preparation.

### Cryo-EM sample preparation

To prepare cryo-EM grids, 2 µL of sample was applied to a freshly glow-discharged (PELCO easiGlow, settings: 0.39 mBar, 25 mA, 25 second glow, 10 second hold) Holey Carbon 300 mesh R 1.2/1.3 gold grid (Quantifoil, Großlöbichau, Germany). After a ∼5 second manual wait time, the grid was blotted (Whatman #1 filter paper) for 3 seconds at blot force 1 and immediately plunge frozen in liquid nitrogen-cooled liquid ethane using a Vitrobot Mark IV (Thermo Fisher Scientific) operated at 4°C and 100% humidity. Grids were clipped after freezing. For grid preparation, the operator was wearing a mask.

### Cryo-EM data collection

All datasets were collected on a Titan Krios G3i electron microscope (Thermo Fisher) operated at 300 kV and equipped with a Gatan BioQuantum Imaging Filter with a slit width of 20 eV. Dose-fractionated images (∼50 electrons per Å^2^ applied over 50 frames) were recorded on a K3 direct electron detector (Gatan) with a pixel size of 1.048 Å. 242 movies were collected around a central hole position using image shift with an 11 x 11 hole pattern and two positions were targeted per hole. The defocus target was varied from −0.6 to −1.6 µm using SerialEM^23^.

### Cryo-EM data processing

9,730 movie stacks were collected, motion-corrected using MotionCor2^24^ in RELION3.1^25^, and CTF-corrected using CTFFIND 4.1.14^26^ (Extended Data Figures 2 – 3 provide additional processing details). Micrographs with a CTFFIND reported resolution estimate greater than 4 Å were discarded, leaving 8,695 micrographs for further processing. An initial particle set of 579 particles was generated by manual picking in RELION and used to train Topaz^27^ in RELION, which in turn was used to pick a set of 20,614 particles from 500 micrographs. These particles were cleaned to 6,531 particles using 2D classification and ab-initio jobs in cryoSPARC^28^. Particles were transferred back to RELION using UCSF pyem tools^29^ and used to train Topaz to pick particles from all 8,695 micrographs, which were then cleaned in cryoSPARC using 2D classification, ab initio, and heterogenous refinement. The process of all-micrograph Topaz particle picking followed by particle cleaning in cryoSPARC was repeated iteratively for three total times, after which the three particle stacks were merged and de-duplicated in RELION with a 100 Å minimum inter-particle distance to generate a stack of 220,293 particles. These particles were non-uniform refined in cryoSPARC4.2.1^30^, then transferred to RELION for post-processing and Bayesian polishing^31^. The polished particle stack was then further cleaned using 2D classification and heterogeneous refinement jobs in cryoSPARC to generate a final “consensus” particle stack of 120,274 particles. The consensus particle stack was non-uniform refined, then iteratively CTF refined (local, then global), followed by a final non-uniform refinement to give a 3.12 Å consensus map which displayed marked heterogeneity in some subunits’ linker regions and LRR domains.

To resolve this heterogeneity, we used 3D variability analysis and display jobs combined with heterogeneous refinement in cryoSPARC^15^. First, using the consensus map, we generated a mask to exclude micellar density. We next applied this mask and the consensus particle stack to a four-component 3D variability analysis job with a 5 Å filter resolution. Next, using the 3D variability display job on simple mode (20 frames, 5 Å filter resolution), we identified two 3D variability components with pronounced conformational changes in subunit linker regions and LRR domains. We used the end volumes (i.e. frames 0 and 19) of these two components produced from the 3D variability display output, alongside a 5 Å low pass-filtered consensus map, as volume references for heterogeneous refinement of the consensus particle stack. Each class was then non-uniform refined, resulting in two well-resolved LRRC8A_BRIL_:D classes (class 0: 3.47 Å, 27,322 particles; class 1: 3.50 Å, 29,827 particles), one class with less resolved LRRs (class 2, 3.48 Å, 26,507 particles), and two classes of significantly lower resolution (class 3: 8.57 Å, 19,744 particles; class 4: 8.44 Å, 16,874 particles). The particles from classes 2 – 4 were pooled together and applied to a second round of heterogenous refinement, with the non-uniformed volumes from classes 0, 1, and 2 as reference volumes. Particles which sorted into classes 0 and 1 in the second round of heterogenous refinement were combined with the initial class 0 and 1 particle stacks and non-uniformed, giving rise to maps for two conformations of LRRC8A_BRIL_:D: conformation 1 (3.29 Å, 43,907 particles, from class 0 particle stacks) and conformation 2 (3.35 Å, 46,893 particles, from class 1 particle stacks). A particle box size of 416 pixels was used throughout and *C*1 symmetry was applied for all 3D jobs.

### Structure modeling, refinement, and analysis

LRRC8A and LRRC8D subunit identity was inferred from the positions of globular densities corresponding to inserted BRIL domains and confirmed with high-resolution features during modeling (Extended Data Figure 4). Sharpened and unsharpened maps from cryoSPARC non-uniform refinement were used to build models in Coot^32^. As a starting point for LRRC8A:D conformation 1, one LRRC8A subunit from a structural model of LRRC8A_BRIL_:C (PDB: 8DS3, chain B)^8^ was trimmed (removing residues 412 – 808) and docked into the LRRC8A:D map for each of the six subunits. For the two LRRC8D subunits, the sequence was then mutated. For LRRC8A:D conformation 2, the LRRC8A:D conformation 1 model was used as a starting point. Models were real-space refined in Phenix^33^ using sharpened maps and assessed for proper stereochemistry and geometry using MolProbity^34^.

The final LRRC8A_BRIL_:D models consist of 1,873 and 1,880 amino acid residues for conformations 1 and 2, respectively. Unmodeled regions consist of each subunit’s N-terminus (residues 1 – 14), extracellular loop 1 (residues 61 – 92, 69 – 92, or 69 – 93 for LRRC8A subunits, along with their inserted BRIL:α-BRIL Fab:α-Fab Nb complexes, residues 61 – 133 for LRRC8D subunits), intracellular loop (residues 175 – 231 for LRRC8A subunits, residues 217 – 276 for LRRC8D subunits), and LRR domain (residues 412 – 810 for LRRC8A subunits, residues 457 – 859 for LRRC8D subunits). In both conformations, 18 lipid molecules were modeled as DOPE (3/18 modeled completely), with ligand restraints obtained from the REFMAC monomer library using Coot.

For display purposes, we docked models for the best resolved LRRs in unsharpened maps using ChimeraX^35^. For LRRC8A subunits, we docked a model of a tightly interacting pair of LRRC8A LRRs obtained from a structural model of LRRC8A_BRIL_:C (PDB: 8DS3, Chains B and C, residues 412 – 808)^8^. For LRRC8D subunits, we used the AlphaFold 3 Server^36^ to predict a model of a LRRC8D monomer which was trimmed to include residues 457 – 848 and docked. Two LRRC8A and two LRRC8D LRRs were docked for conformation 1, four LRRC8A LRRs were docked for conformation 2.

Pore measurements were made using HOLE^37^. Figures were prepared using ChimeraX, Prism, and Adobe Illustrator software.

### Electrophysiology

HeLa *LRRC8A-E^−/–^* cells^19^ were cultured in DMEM (Gibco) supplemented with 10% FBS (Gibco), 100 units/mL penicillin and 100 μg/mL streptomycin (Gibco) and grown at 37 °C in the presence of 5% CO_2_. Trypsinized cells were deposited on 5 mm glass coverslips in a 6-well dish six hours to two days prior to transfection. At least 30 minutes prior to transfection, wells were replaced with antibiotic-free growth media. Consistent with a prior report^38^, we were unable to reproducibly obtain large wild-type VRAC currents with a 1:1 transfection ratio of LRRC8A and LRRC8D plasmids. Thus, for all transfections, a 1:5 mass ratio of LRRC8A and LRRC8D plasmids was used and mixed with FuGENE 6 at a 1:3 volume ratio diluted in serum- and antibiotic-free DMEM (typically, 0.5 μg LRRC8A and 2.5 μg LRRC8D mixed with 9 μL FuGENE 6 in 100 μL DMEM). Media was replaced 12 – 24 hours post-transfection with growth media containing antibiotics. Patch clamp experiments were conducted 36 – 48 hours post-transfection.

For patch clamp recording, coverslips were placed in a perfusion chamber at room temperature in isotonic bath solution (90 mM NaCl, 2 mM KCl, 1 mM CaCl_2_, 1 mM MgCl_2_, 10 mM HEPES, 10 mM glucose, adjusted to pH 7.4 with NaOH). Mannitol was used to adjust the solution osmolarity to ∼330 mOsm (the approximate osmolarity of the cell culture media) as measured by a vapor pressure osmometer (VAPRO, Model 5600, ELITechGroup) or a freezing-point depression osmometer (Model 3320, Advanced Instruments). Borosilicate glass pipettes were pulled to a resistance of ∼1.5 – 4 MΩ and filled with pipet solution (133 mM CsCl, 5 mM EGTA, 2 mM CaCl_2_, 1 mM MgCl_2_, 10 mM HEPES, ∼4 mM MgATP, adjusted to pH 7.4 with CsOH and ∼330 mOsm with mannitol). An Axopatch 200B amplifier connected to a Digidata 1550B digitizer (Molecular Devices) was used for data acquisition with pClamp10.7 software. Analog signals were filtered at 1 kHz and sampled at 10 kHz. Pressure application from the patch pipette was accomplished with a high-speed pressure clamp (HSPC, ALA Scientific). Once a patch was achieved in the whole cell mode and a stable whole-cell capacitance was measured, voltage families were recorded to monitor pre-swelling currents with the following voltage protocol applied every 46 s: V_hold_ = 0 mV; V_test_ = −100 to +100 mV, Δ20 mV, 400 ms. When initial currents stabilized, hypotonic bath solution (same components as isotonic, but adjusted to ∼250 mOsm with mannitol) was exchanged into the chamber and currents were monitored over the course of cell swelling until currents stopped increasing for multiple records or the patch broke. If a patch exhibited signs of leaking or sealing, a membrane test protocol was examined to either reseal, re-break-in, or discard the patch. Care was taken to make sure that the measured pressure was slightly negative (i.e., −1 mm Hg) after the whole-cell mode was achieved. Cells were monitored using a brightfield microscope and were selected for patching based on cell morphology (aiming for healthy interphase cells) and the presence of fluorescent reporters (GFP and mCherry).

Data was analyzed using Clampfit 10.7, Excel, and Graphpad Prism 10 software. To quantify current densities, the mean current density from the last 50 ms of each trace prior to the onset of capacitive currents was measured using Clampfit and normalized by the cell’s measured capacitance. The displayed data are current densities at 60 mV (where VRAC current inactivation is minimal) averaged from three consecutive records after achieving stable currents in isotonic or hypotonic solution and the ratio of these current densities was used to measure fold activation. For each cell, the reversal potential was measured in Prism as the x-intercept (voltage) of a linear fit of the current amplitudes at 0 mV and −20 mV from three averaged consecutive hypotonic records. For plotting representative current traces, a single isotonic and hypotonic record was chosen, the data was decimated 10-fold (to 1 ms sampling interval) and plotted in Prism. Prism was used for statistical analyses of fold activation. A Brown-Forsythe and Welch one-way ANOVA test followed by a Dunnett’s T3 multiple comparisons test was used to compare all mutant channels to wild-type LRRC8A:D. An unpaired *t* test was used to compare LRRC8A(T48D):D and LRRC8A(T48D):D(T48D).

#### Acknowledgements

We thank Dan Toso, Ravindra Thakkar, and Paul Tobias for microscope and computational support at the Cal-Cryo facility. We thank the laboratory of Ardem Patapoutian for generously providing the *LRRC8* knock-out HeLa cell line. We thank the laboratory of Anthony Kossiakoff for providing the α-BRIL Fab BAG2 and α-Fab Nb. We thank members of the Brohawn laboratory for discussions and feedback on the manuscript. This work was funded by The New York Stem Cell Foundation, a McKnight Foundation Scholar Award, and a Sloan Research Fellowship to S.G.B., NIGMS grant no. GM128263 to D.M.K., and a Shurl and Kay Curci Ph.D. Scholarship to A.L.

## Contributions

D.M.K. designed the fiducial tagging approach. D.M.K. and K.H. performed biochemical experiments and collected cryo-EM data. A.L. processed cryo-EM data, modeled and refined the structures, generated mutant constructs, performed electrophysiology experiments, and analyzed all presented data. A.L., D.M.K., and S.G.B. conceived of the project. A.L. and S.G.B. wrote the manuscript with input from all authors.

## Data availability

For LRRC8A_BRIL_:D conformation 1, the final model is deposited in the PDB under 9DX7 and the final map is deposited in the EMDB under EMD-47282. For LRRC8A_BRIL_:D conformation 2, the final model is deposited in the PDB under 9DXA and the final map is deposited in the EMDB under EMD-47283. Original micrograph movies and final particle stacks are deposited in the Electron Microscopy Public Image Archive (EMPIAR) under [*to be inserted upon deposition*].

---

**Extended Data Table 1.**
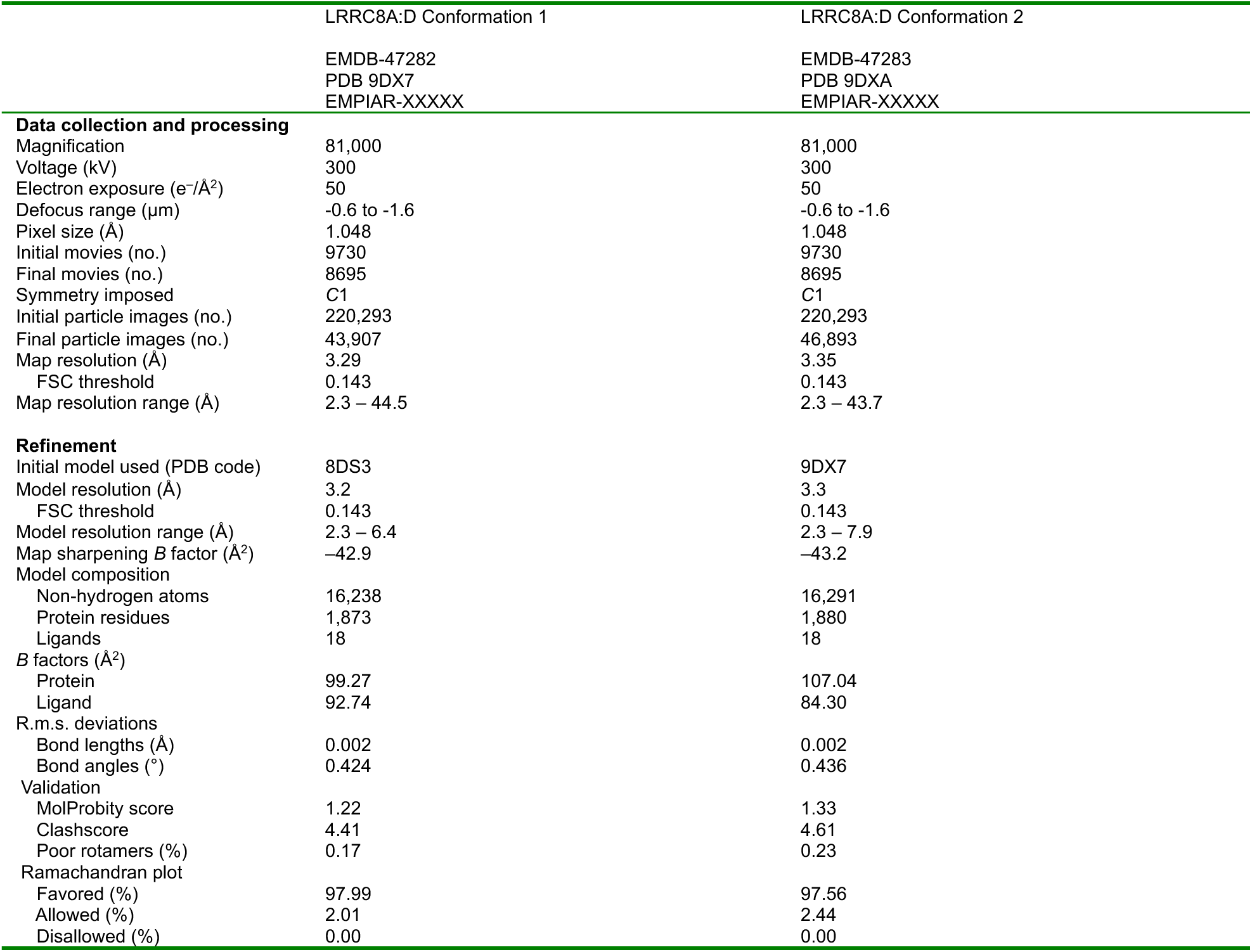
Cryo-EM data collection, refinement and validation statistics.

